# Causal evidence for a cerebellar role in prefrontal-hippocampal interaction in spatial working memory decision-making

**DOI:** 10.1101/2020.03.16.994541

**Authors:** Yu Liu, Samuel S. McAfee, Meike E. Van Der Heijden, Mukesh Dhamala, Roy V. Sillitoe, Detlef H. Heck

**Affiliations:** Department of Anatomy and Neurobiology, University of Tennessee HSC, Memphis, TN; Department of Diagnostic Imaging, St Jude Children’s Research Hospital, Memphis, TN; Department of Neuroscience, Baylor College of Medicine; Department of Physics and Astronomy, Neuroscience Institute, Georgia State University; Department of Pathology and Immunology, Baylor College of Medicine; Development, Disease Models & Therapeutics Graduate Program, Baylor College of Medicine; Jan and Dan Duncan Neurological Research Institute of Texas Children’s Hospital, Houston, TX

**Keywords:** Cerebellum, Purkinje cell, coherence, spatial working memory, hippocampus, medial prefrontal cortex, cognition, electrophysiology, optogenetics

## Abstract

Spatial working memory (SWM) is a cognitive skill supporting survival-relevant behaviors, such as optimizing foraging behavior by remembering recent routes and visited sites. It is known that SWM decision-making in rodents requires the medial prefrontal cortex (mPFC) and dorsal hippocampus. The decision process in SWM tasks carries a specific electrophysiological signature of a brief, decision-related increase in neuronal communication in the form of an increase in the coherence of neuronal theta oscillations (4-12 Hz) between the mPFC and dorsal hippocampus, a finding we replicated here during spontaneous exploration of a plus maze in freely moving mice. We further evaluated SWM decision-related coherence changes within frequency bands above theta. Decision-related coherence increases occurred in seven frequency bands between 4 and 200 Hz and decision-outcome related differences in coherence modulation occurred within the beta and gamma frequency bands and in higher frequency oscillations up to 130 Hz. With recent evidence that Purkinje cells in the cerebellar lobulus simplex (LS) represent information about the phase and phase differences of gamma oscillations in the mPFC and dorsal hippocampus, we hypothesized that LS might be involved in the modulation of mPFC-hippocampal gamma coherence. We show that optical stimulation of LS significantly impairs SWM performance and decision-related mPFC-dCA1 coherence modulation, providing causal evidence for an involvement of cerebellar LS in SWM decision making at the behavioral and neuronal level. Our findings suggest that the cerebellum might contribute to SWM decision-making by optimizing the decision-related modulation of mPFC-dCA1 coherence.

## INTRODUCTION

Neural representations of the environment and the ability to form a spatial memory is a survival-relevant function, allowing complex behaviors such as foraging for food, building nests, marking territories, and finding escape routes. Each of these behaviors relies in part on spatial working memory (SWM) to optimize decisions about which path or target location to choose or avoid based on recent experiences (Amemiya and Redish, 2018; Eichenbaum, 2017). In rodents, decision-making in SWM requires the coordinated activity of the medial prefrontal cortex (mPFC) and dorsal hippocampus (Churchwell and Kesner, 2011; Gordon, 2011). Electrophysiological recordings in rodents during performance of SWM tasks involving some form of rule learning have shown that the decision process is associated with an increase in the coherence of theta oscillations between the mPFC and dorsal hippocampus (Benchenane et al., 2011; Gordon, 2011; Jones and Wilson, 2005) and a concomitant increase in entrainment of mPFC spike activity to the phase of hippocampal theta oscillations (Hyman et al., 2010; Jones and Wilson, 2005). Two studies compared theta coherence changes between correct and incorrect decisions and found that the magnitude of coherence increase reflected decision outcome. Theta coherence between the mPFC and dorsal hippocampus reached higher values during correct compared to incorrect decisions (Hyman *et al*., 2010; Jones and Wilson, 2005). The modulation of coherence, particularly coherence of gamma (~ 40 Hz) oscillations has been proposed as a mechanism for the precise spatial and temporal coordination of neuronal communication between cerebral cortical areas during cognitive processes (Fries, 2005; Womelsdorf et al., 2007). The proposed principle of “communication through coherence” has since received substantial support from experimental findings showing that increased coherence does affect spike activity in communicating structures by increasing phase-locking of spike activity and increasing spike synchrony (Bosman et al., 2012; Brunet et al., 2014; Jones and Wilson, 2005; McAfee et al., 2018; Siegel et al., 2008).

Correlational evidence has implicated the cerebellum in working memory tasks involving visual and verbal working memory in humans (Desmond et al., 1997; King et al., 2019) and spatial memory in mice (Badura et al., 2018). Known multi-synaptic pathways connect the cerebellum with the two cerebral cortical structures that are essential for SWM, the mPFC (Kelly and Strick, 2003; Middleton and Strick, 2001) and the dorsal hippocampus (Watson et al., 2019).

We recently showed that Purkinje cells in the cerebellar lobulus simplex (LS) represent phase information of gamma oscillations in the mPFC and dorsal hippocampal CA1 region (dCA1) (McAfee et al., 2019), leading us to hypothesize that LS Purkinje cells might be involved in modulating decision-related mPFC-dCA1 coherence. Here, we use a combination of *in vivo* electrophysiological recordings in the mPFC and dCA1 and optogenetic activation of cerebellar Purkinje cells in LS in freely moving mice to investigate the role of the LS in SWM decision-making and the modulation of mPFC-dCA1 coherence during spontaneous exploration of a plus maze.

## RESULTS

Experiments were conducted in 9 adult transgenic mice (4 female, 5 male), co-expressing Channelrhodopsin2 fused to an enhanced yellow fluorescent protein (EYFP) in cerebellar Purkinje cells Tg(Pcp2-COP4*H134R/EYFP) U126Isop/J). We quantified SWM performance by evaluating the arm choices mice made during 12 min of spontaneous exploration of a plus maze (Fig. 1). During such spontaneous maze explorations, healthy mice have an instinctive tendency to avoid re-entering recently visited maze arms, which makes it possible to measure SWM performance by extracting the percentage of spontaneous alternations from the sequence of arm entries (Lalonde, 2002; Richman et al., 1987). Spontaneous alternations are defined as sequences of four consecutive arm entries without repeat (Fig. 1B) and the percentage of spontaneous alternations out of the entire sequence of entries serves as quantitative measure of SWM (Lalonde, 2002; Richman et al., 1987). Chance performance in a four-arm maze corresponds to 22.2% spontaneous alternations. Healthy mice generate spontaneous alternations significantly above chance level (Richman *et al*., 1987)

**Figure 1.**
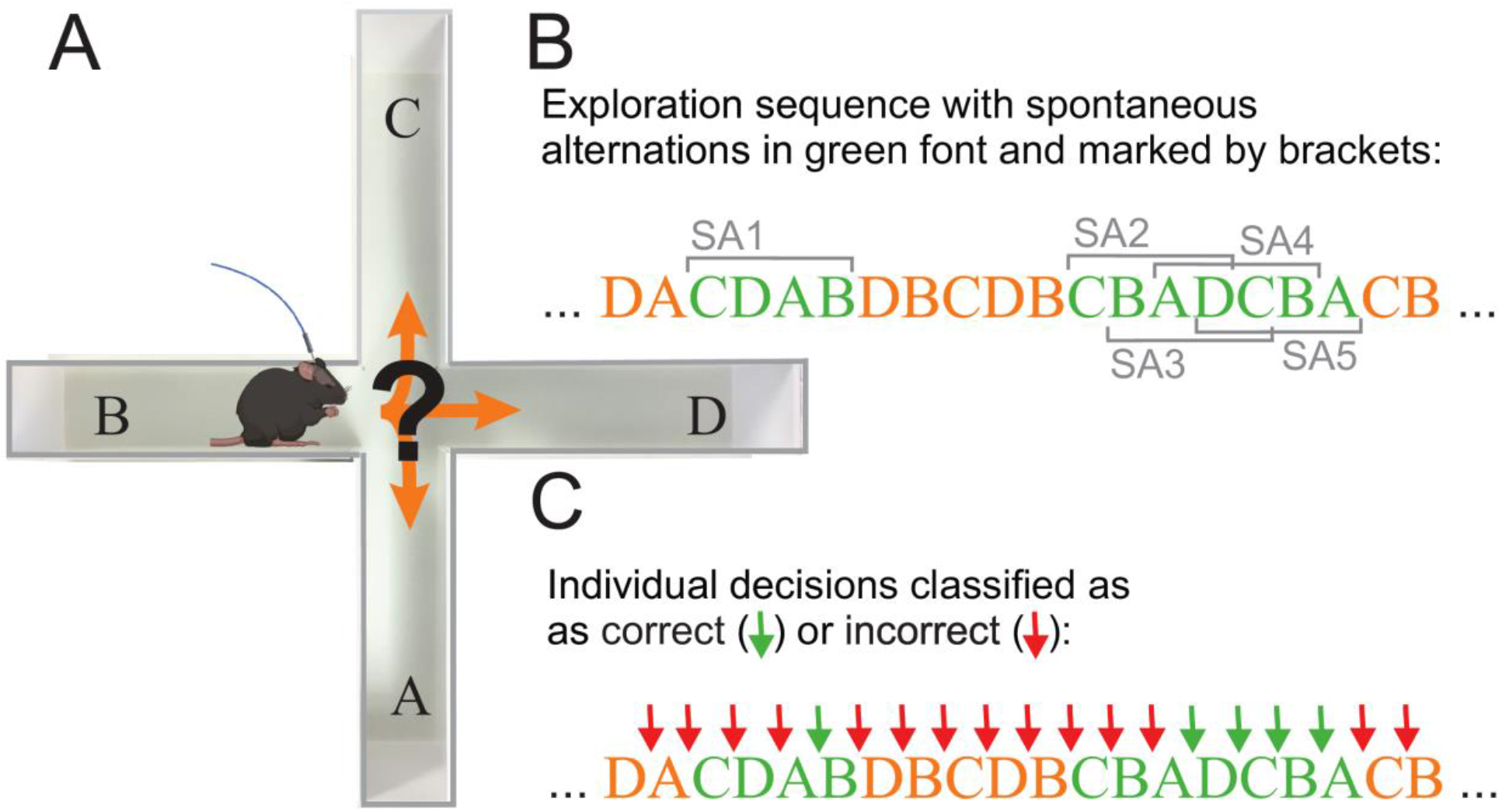
Evaluation of plus-maze exploration behavior, spatial working memory and classification of individual decisions. **(A)** Mice implanted with recording electrodes in the mPFC and dCA1 and an optical fiber above cerebellar LS were placed in a plus maze and allowed to freely explore for 12 min. Movement trajectories were video-captured and digitized for offline analysis. **(B)** The sequence of arm entries was analyzed to detect any sequence of four entries without a repeat (i.e. spontaneous alternations). In the example shown arm entries that are part of a spontaneous alternation are printed in green font. A total of five spontaneous alternations are marked with grey brackets labeled SA1-5. Note that spontaneous alternations can be overlapping (SA2-5). The first occurrence of a spontaneous alternation in the example is a single sequence of four entries (CDAB, SA1). The second occurrence (CBADCBA) contains four overlapping spontaneous alternations (CBAD, BADC, ADCB, DCBA, SA2-5). **(C)** Same sequence as in (B) but with individual arm entry decisions classified as correct (green arrows) or incorrect (red arrows) based on the position of the decision in a sequence of four entries. For the analysis of decision-outcome related neuronal activity only decisions that were preceded by three choices without repetition were classified as correct. As a consequence, the first two or three decisions of a spontaneous alternation (CBAD) were classified as incorrect depending on the choice immediately preceding the spontaneous alternation.

In order to link brain activity to decision-making outcome, we classified individual decisions as correct or incorrect, depending on whether the decision completed a spontaneous alternation (Fig. 1C) (see methods for details).

Mice had two extracellular recording electrodes (glass insulated tungsten/platinum; 80 μm outer diameter; impedance: 3.5–5.0 MΩ, Thomas Recording, Germany) implanted into each of their mPFC and dCA1 (Fig. 2A). An LED light source (465 nm, 17 mW, Doric Lenses, Canada) attached to an optical fiber (200 μm, Doric Lenses, Canada) was mounted over the LS with the fiber tip touching the intact dura (Fig. 2). Recording electrodes captured local field potentials (LFPs) in the mPFC and dCA1 (Fig. 2B). Correct electrode placement in dCA1 was verified using the presence of sharp wave ripples in the signals (Fig. 2B). From the two recording sites in each structure, only one was chosen for further analysis. If there was a difference in noise levels, the less noisy signal was chosen. All recording sites were marked with electrolytic lesions and thereafter verified anatomically (Fig. 2C).

**Figure 2.**
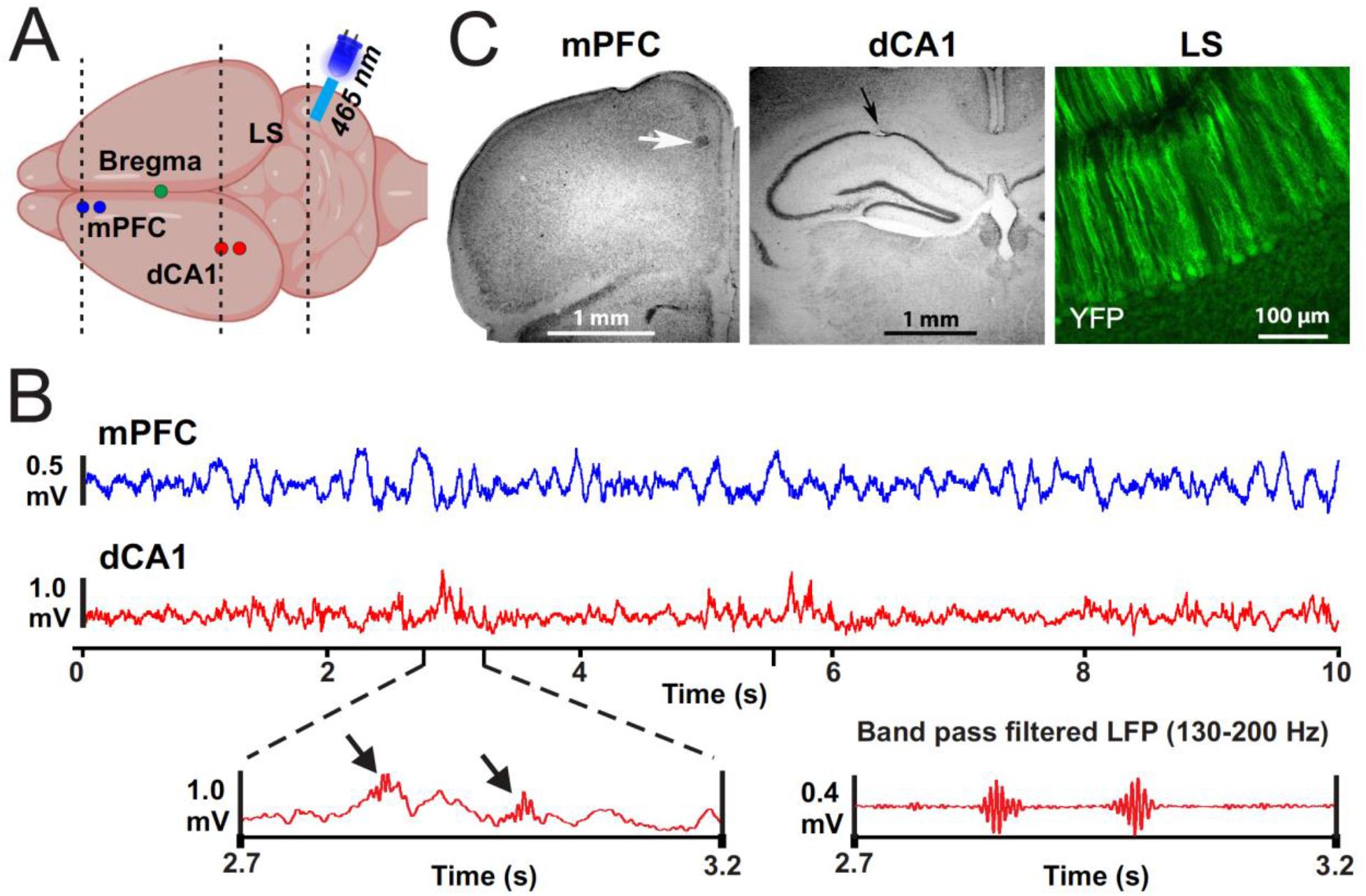
Illustration of recording locations, example data, and lesion sites. **(A)** Schematic drawing of the top view of a mouse brain with recording locations in the left mPFC and dCA1 marked with blue and red circles respectively. The illustration above the cerebellum depicts an optical fiber coupled to an LED (465 nm) that was mounted over LS. Dashed lines indicate the approximate locations of coronal sections shown in panel (C). **(B)** Top traces are LFP signals recorded in the mPFC (blue) and dCA1 (red). The left bottom panel shows an enlarged sections of the dCA1 LFP that includes sharp-wave ripple activity indicated by black arrows. The bottom right panel shows a band pass filtered (130-200Hz) version of the LFP on the left, highlighting the two sharp wave ripple events. **(C)** Examples of electrolytic lesions (arrows) at recording sites in Nissl-stained sections of the mPFC and dCA1 region and a coronal section of cerebellar LS showing YFP expression in PCs of the L7-ChR2-GFP mice.

### Optogenetic activation of the LS impairs SWM performance and decision-related LFP activity

Plus-maze spontaneous exploration sessions with and without light stimulation were randomized. During sessions with LS stimulation, a 1 sec light stimulus (sinusoidal modulation at 120 Hz) was manually timed to occur as the mouse approached the center square of the maze, defined by the mouses head having entered the blue shaded area in Fig. 3A, right panel. Mouse behavior was not visibly affected by the stimulus and as a result the mice continued to move towards the center of the maze and the next arm in the same fashion as during control trials. Comparing SWM performance between control and LS stimulated trials showed a significant reduction of spontaneous alternations. During control trials mice generated 38.3% spontaneous alternations, which is significantly more than expected by chance (p=0.0004, Two-Way ANOVA). During LS stimulation trials mice generated 24.2%, which was similar to chance level performance (p>0.8, Two-Way ANOVA) and significantly below control trial performance (p=0.0012, Two-Way ANOVA) (Fig. 3B).

**Figure 3.**
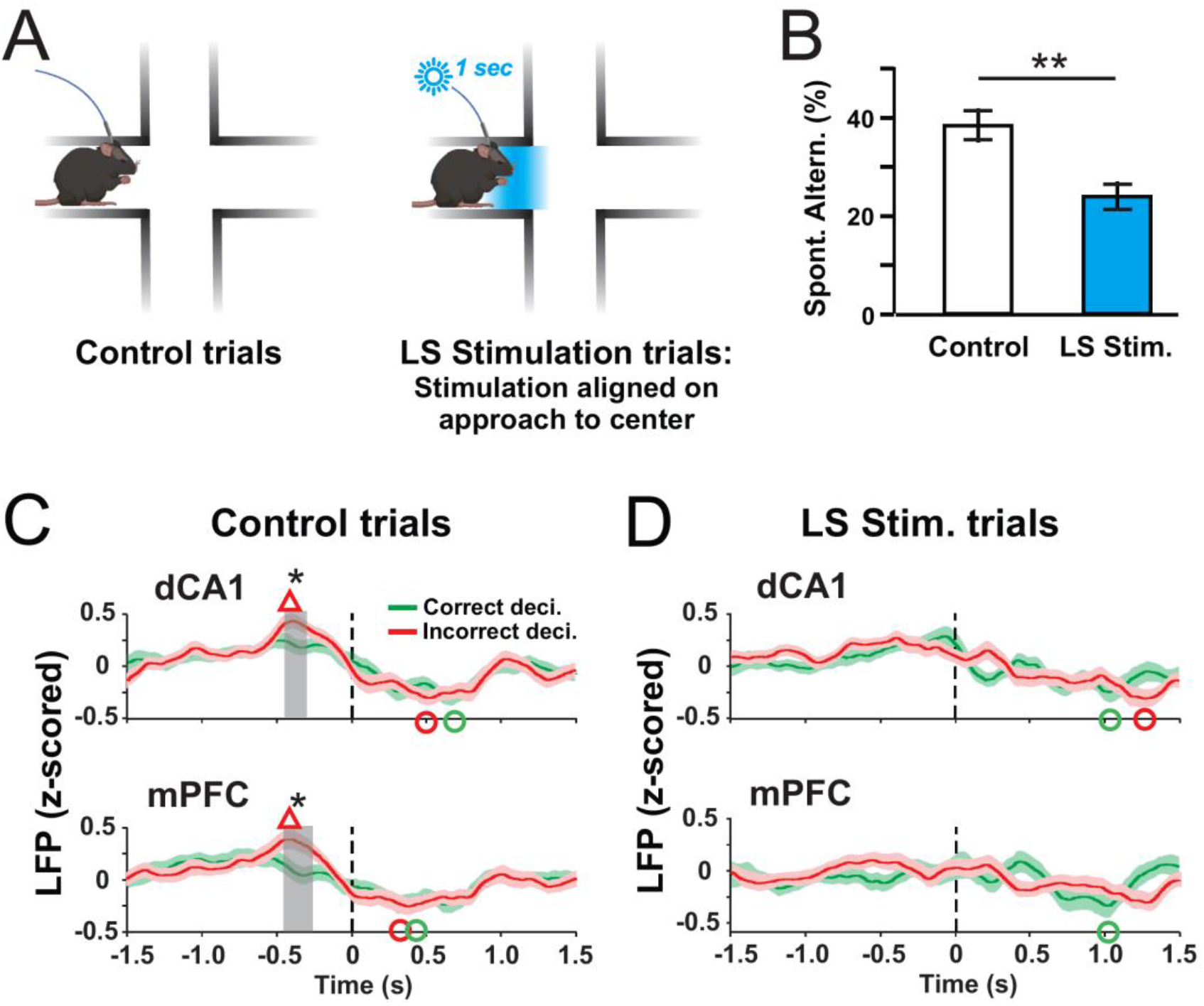
Optogenetic activation of PCs in the LS impairs SWM performance and decision-outcome related modulation of LFP activity. **(A)** During control trials mice freely explored the maze for 12 min without optical stimulation (left panel). During trials with LS optical stimulation (right panel) LS PCs were photo-activated for 1 sec at the time the mouse reached the center of the maze. Photo activation was started manually when the mouse’s head reached the area marked in blue. **(B)** Compared to control trials, the percentage of spontaneous alternations was significantly reduced in LS stimulation trials (**P = 0.0012; Two Way ANOVA). **(C)** Average time courses of decision-related LFP potentials in the dCA1 and mPFC for correct (green traces) and incorrect (red traces) decisions. Time zero in each plot corresponds to the time the mouse left the maze center with all four paws. Average peri-decision LFP activities in both dCA1 and mPFC reached values at around −0.4 sec that were higher during incorrect than correct decisions. Red triangles mark peak LFP values for incorrect decisions that exceeded baseline values calculated by averaging pre-decision (−1.5 to −1.25 s) values (P<0.05; Wilcoxon Signed-Rank test). Green and red circles mark LFP minima that fall below pre-decision average for correct and incorrect decisions, respectively (P<0.05; Wilcoxon Signed-Rank test). Grey shaded rectangles mark time periods with significant differences between the traces (*P<0.05; Wilcoxon Signed-Rank test). **(D)** As in (C) but for LS stimulation trials. LS stimulation eliminated the decision-outcome related differences in dCA1 and mPFC LFP activity that preceded decision completion (~ −0.4 sec) during control trials.

LFP activities in the mPFC and dCA1 were continually recorded during the 12 min of exploration. For further analysis, 3 seconds of peri-decision LFP activity was extracted, aligned on the time of completion of the decision process defined as the time when the mouse had left the maze center with all four paws (t = 0 sec in all plots of Fig. 3C,D). We evaluated the peri-decision LFP activity changes relative to pre-decision baseline values calculated as the average LFP activity in the −1.5 to −1.25 s time period, relative to the time of decision completion at t = 0 (Fig. 3). In control trials, LFP activity reached above baseline peak values (marked with red and green triangles) in both dCA1 (peak time −0.39 s; p=0.000) and mPFC (peak time −0.43 s; p=0.007) when mice made incorrect decisions (Fig. 3C). LFP activities in both the mPFC and dCA1 reached below baseline minima (marked with red and green circles) after the decisions were completed for both correct (trough time: mPFC 0.31s, p<0.0001; dCA1 0.67 s, p<0.0001) and incorrect decisions (trough time: mPFC 0.38 s, p=0.0003; dCA1 0.51s, p=0.001).

During control trials, peak LFP activity in the mPFC and dCA1 differed between correct and incorrect trials. At around 0.4 s prior to completion of the decision process, LFP values measured during correct decisions were significantly lower than values measured during incorrect decisions (grey shaded areas, dCA1 −0.42 to −0.32 s, mPFC −0.44 to −0.28 s; *p* < 0.05; Two-Sample t-test) (Fig. 3C). During LS stimulation trials, by contrast, the decision-outcome related differences in LFP activity were no longer observed in neither mPFC nor dCA1. There were no significant differences between the correct and incorrect decisions in the LFP minima (Fig. 3C).

### Peri-decision maxima and minima in mPFC-dCA1 coherence

We conducted a time-resolved analysis of coherence of mPFC-dCA1 LFP activity for neuronal oscillations between 4 and 200 Hz divided into eight commonly used frequency bands: theta (4-8 Hz), alpha (8-12 Hz), beta (12-30 Hz), low gamma (30-60 Hz), mid gamma (60-80 Hz), high gamma (80-100 Hz), high frequency (100-130 Hz) and very high frequency (130-200 Hz). Results were separated by decision-outcome to compare peri-decision time courses of mPFC-dCA1 coherence for correct and incorrect decisions (Fig. 4). As described above for the analysis of peri-decision LFP activity, time resolved coherence results were aligned on the time of completion of the decision process (t = 0 in all plots in Fig. 4). In order to allow a comparison of coherence modulation across mice, baseline coherence values were subtracted from each point of the peri-decision coherence function, with baseline coherence defined as the average coherence in the 0.5 s period immediately after the mouse entered the center square of the maze with all for paws (see Methods).

**Figure 4:**
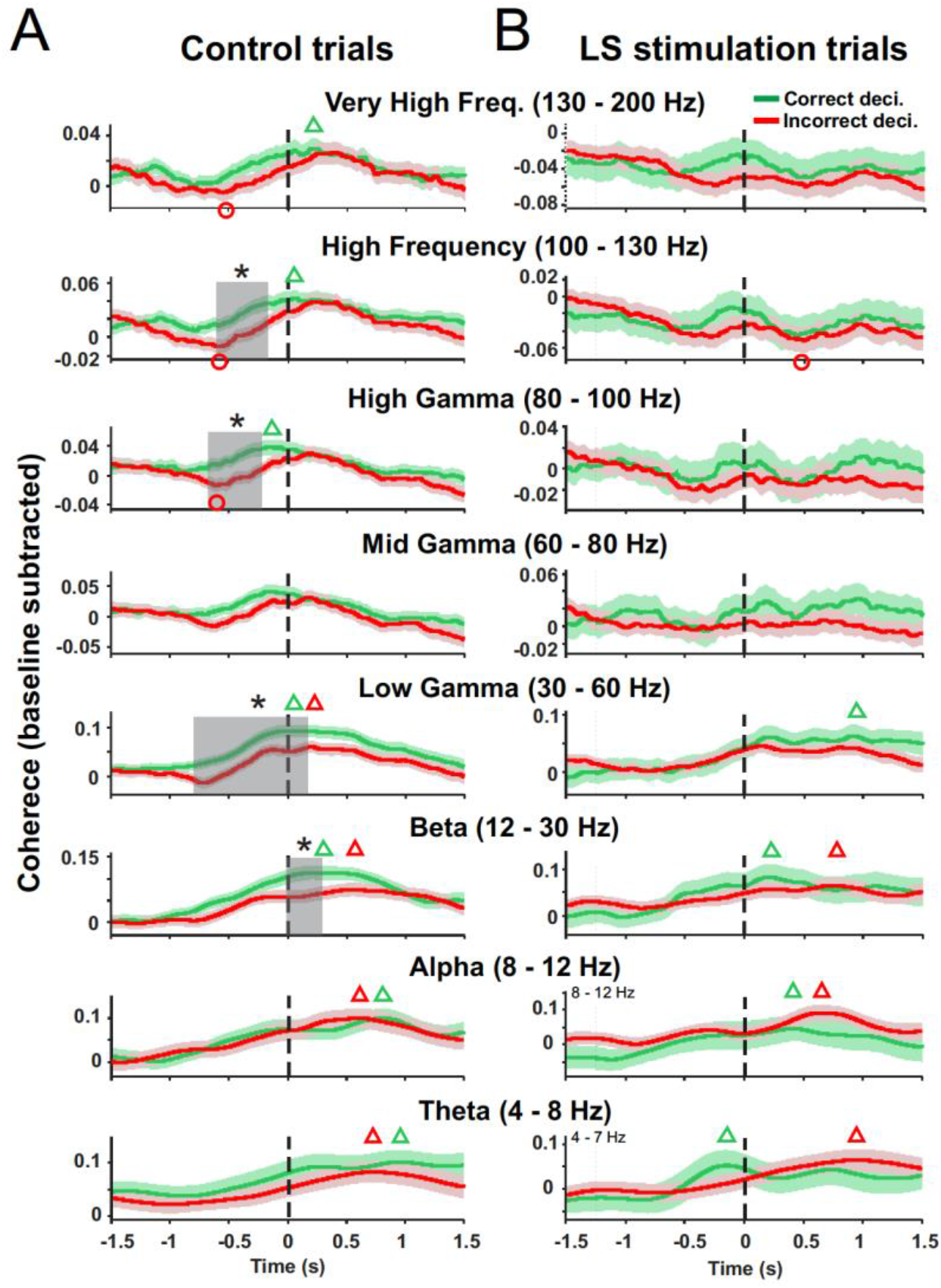
Optical stimulation of LS Purkinje cells disrupts decision-related mPFC-dCA coherence modulation. **(A)** Time resolved peri-decision coherence averages for correct (green traces) and incorrect (red traces) decisions for control trials. Peri-decision coherence was analyzed for eight separate frequency bands between 4 and 200Hz with time 0 making the completion of the decision process defined by the mouse entering the next maze arm with all four paws. Baseline coherence values for each mouse were subtracted from peri-decision coherence functions before averaging. Green and red triangles mark coherence peaks for correct and incorrect decisions, respectively, that exceed baseline coherence (Wilcoxon Signed-Rank test P<0.05). Red circles mark coherence minima falling below baseline coherence values for incorrect decisions (Wilcoxon Signed-Rank test; P<0.05). No significant coherence minima were found for correct decisions. Decision outcome was reflected in differences in time course of coherence functions. Peri-decision times where coherence functions for correct and incorrect decisions differed are marked by grey shaded rectangles (Two-sample t-test, *p < 0.05). **(B)** As in (A) but for LS-stimulation trials. LS stimulation eliminated all differences between peri-decision coherence functions for correct and incorrect decisions.

Next, we asked whether coherence in any frequency band increased or decreased significantly relative to baseline during the decision process and whether these changes were linked to decision outcome. Coherence minima and maxima were detected in the average coherence functions and tested for significance relative to baseline coherence for correct and incorrect decisions using Wilcoxon Signed-Rank test. In Fig. 4 green and red triangles mark significant coherence maxima for correct and incorrect decisions, respectively, and red circles mark significant coherence minima for incorrect decisions. No significant minima were found for correct decisions.

In control trials, peri-decision coherence in the theta, alpha, beta, and low gamma bands reached above-baseline peak values (p<0.05) irrespective of decision outcome, with all peaks occurring after the decision process was completed (t>0, Fig. 4A). Coherence peaks occurred earliest in the gamma frequency band (t=0.02 s correct, t=0.21 s incorrect) and at increasingly later times for lower frequencies: beta (t=0.21 s correct, t=0.56 s incorrect), alpha (t=0.81 s correct, t=0.62 s incorrect), and theta (t=0.96 s correct, t=0.72 s incorrect). In the mid gamma band, peri-decision coherence did not significantly deviate from baseline levels (Fig. 4A). In the high gamma, high frequency and very high frequency bands peri-decision coherence reached above-baseline peak values for correct decisions only, with coherence peaking around the time of decision completion (high gamma t=−0.17 s, high frequency t=0.06 s, very high frequency t=0.22 s, Fig. 4A). By contrast, incorrect decisions were associated with below-baseline coherence minima in the high gamma, high frequency and very high frequency bands which preceded decision completion (Fig. 4A). Coherence minima at similar time points occurred during incorrect decisions in the mid and low gamma ranges trending towards statistical significance (mid gamma: t=-0.62 s; p=0.0504; low gamma: t=−0.72 s; p=0.0690).

### Optogenetic activation of the LS impairs decision-outcome related mPFC-dCA1 coherence

In control trials, the time resolved coherence functions of beta (12-30 Hz), low gamma (30-60 Hz), high gamma (80-100 Hz) and high frequency (100-130 Hz) oscillations differed significantly between correct and incorrect decisions (Two-sample t-test, p<0.05), with correct decisions associated with higher coherence values during periods of significant coherence differences (grey shaded areas Fig. 4A). Decision related differences in coherence occurred before decision completion in the high gamma and high frequency bands and right after decision completion in the beta band (Fig. 4A). In the low gamma band, decision-related coherence differences occurred during an extended period starting at 0.80 s prior to decision completion and ending at 0.11 s after completion (Fig. 4A). No differences in decision-outcome related coherence were observed in frequencies bands below 12 Hz or above 130 Hz. Stimulation of LS Purkinje cells eliminated decision-related differences in mPFC-dCA1 coherence in all frequency bands (Fig. 4B).

In LS-stimulation trials peri-decision coherence in the theta, alpha, and beta bands reached above-baseline peak values (p<0.05) irrespective of decision outcome (Fig. 4B). The timing of coherence peaks relative to decision-completion was similar to peak timing in control trials, with the exception of the theta band, where coherence peaked prior to decision completion in LS-stimulation trials compared to after decision completion in control trials (compare Figs. 4 A and B). In contrast to control, trials peak coherence in the low gamma band no longer reached above-baseline level values for incorrect decisions while above-baseline coherence peaked at about 0.9 sec after decision-completion compared to 0.02s in control trials (Fig. 4A,B). Compared to control trials, LS–stimulation prevented correct-decision-associated coherence increases above baseline in the high gamma, high frequency and very high frequency bands (Fig. 4A,B). Below-baseline coherence minima, which occurred prior to decision completion in the high-gamma, high-frequency and very-high frequency bands in control trials, were eliminated in LS–stimulation trials (Fig. 4A,B). A below-baseline coherence minimum associated with incorrect decisions during LS-stimulation trials was observed in the high frequency bands at 0.5 s after decision completion.

## DISCUSSION

We investigated the involvement of the mouse cerebellum in SWM decision-making by quantifying spontaneous alternations during free exploration of a plus maze (Fig. 1). We used extracellular recording techniques to measure decision-related neuronal activity in the mPFC and dCA1 and optogenetic tools to modulate neuronal activity in cerebellar LS during the decision-making process. Our findings provide causal evidence for a cerebellar role in SWM decision-making at the level of both behavior and neuronal activity. In addition, our results expand previous findings of decision-related mPFC-dCA1 theta coherence modulation (Benchenane et al., 2010; Hyman *et al*., 2010; Jones and Wilson, 2005; Sigurdsson et al., 2010) by showing that decision-related coherence modulation occurs in multiple frequency bands between 4 and 200Hz. Previous studies reported decision-outcome related differences in theta coherence modulation (Hyman *et al*., 2010; Jones and Wilson, 2005). We found decision-outcome related differences in coherence modulation in multiple frequency bands, but interestingly not in the theta band. We suggest that the lack of theta coherence correlation with decision outcome is due to the lack of a rule learning requirement in the spontaneous exploration task used here (see below). We also found significant decreases in coherence that preceded coherence increases during incorrect decisions, which adds a new feature to the link between SWM decision outcome and mPFC-dCA1 coherence modulation.

In rodents SWM is known to jointly require the mPFC and dCA1 (Churchwell and Kesner, 2011). The prefrontal cortex is presumed to integrate spatial information encoded in the hippocampus with working memory information about recent pathways and arm choices in order to optimize foraging behavior (Jones and Wilson, 2005). There is substantial evidence that the interaction between the mPFC and hippocampus during SWM based decision-making involves increases in the coherence of theta oscillations (Benchenane *et al*., 2010; Hyman *et al*., 2010; Jones and Wilson, 2005; Sigurdsson *et al*., 2010) and enhanced coupling of spiking activity in the mPFC to the phase of hippocampal theta oscillations (Hyman *et al*., 2010; Jones and Wilson, 2005). Task-specific coherence modulations are believed to modulate functional connectivity by facilitating neuronal information exchange through the proper alignment of time windows of maximal excitability in communicating brain structures (Fries, 2015; Womelsdorf *et al*., 2007). This notion of “communication through coherence” is supported by findings directly relevant to SWM, showing that spike activity in the mPFC becomes increasingly phase locked to hippocampal LFP oscillations during periods of high mPFC-hippocampal theta coherence in a SWM (Jones and Wilson, 2005; Sigurdsson *et al*., 2010) or a spatial delayed-non-match-to-sample task (Hyman *et al*., 2010).

Two previous studies compared mPFC-hippocampal theta coherence based on decision-outcome and showed higher coherence values during correct compared to incorrect decisions (Hyman *et al*., 2010; Jones and Wilson, 2005). Our results confirm that theta coherence increases during decision-making but we did not find differences in theta coherence between correct and incorrect decisions (Fig. 4A) Instead we identified decision-outcome related differences in the coherence of mPFC-dCA1 beta (12-30Hz), low gamma (30-60Hz), high-gamma (80-100Hz) and high frequency (100-130Hz) oscillations. Consistent with previous findings for decision-outcome related theta coherence, in each frequency band correct decisions were associated with higher coherence values than incorrect decisions (Fig. 4A). In addition, we found that in the high gamma, high frequency and very high frequency range incorrect decisions were associated with significant decreases in coherence that preceded coherence increases. Coherence in the low and mid gamma ranges also showed decreases associated with incorrect decisions that trended towards significance. In all five frequency bands that showed coherence decreases during incorrect decisions, coherence minima occurred within a 160 ms time window (0.55-0.71 s) around 0.62 s before completion of the decision process (Fig. 4A). The timing of significant coherence peaks associated with correct decisions occurred close to or after the time of decision completion (Fig. 4A). Just based on their time of occurrence prior to decision completion, differences in decreases in coherence during the decision-process seems more likely to be causally linked to decision outcome than differences in peak coherence values. The coherence decrease may be indicative of failed or sub-optimal information exchange between the mPFC and hippocampus during a critical time during the decision process.

The cerebellum is functionally connected with both the mPFC (Kelly and Strick, 2003; Middleton and Strick, 2001) and the dorsal hippocampus (Watson *et al*., 2019), two anatomical pathways that provide a potential substrate for cerebrocerebellar communication during SWM. Imaging studies in humans have shown activation of the LS / Crus I areas of the cerebellum during visual and verbal working memory tasks (Desmond *et al*., 1997; King *et al*., 2019). We have recently shown that Purkinje cells in the LS and Crus I represent phase information of neuronal oscillations in the mPFC and dCA1 (McAfee *et al*., 2019), with gamma phase information only represented in the LS. This finding which led us to hypothesize that LS Purkinje cells might have a role in modulating decision-related mPFC-dCA1 gamma coherence and thus SWM decision-making. Behavioral and electrophysiological results from this study support our hypothesis. Comparing SWM behavioral performance in terms of percentage of spontaneous alternations shows that stimulation of LS Purkinje cells reduced the number of correct SWM decisions to around chance level, providing causal evidence for the involvement of LS in SWM decision-making (Fig. 3B). Analysis of neuronal activity in control trials showed average peri-decision LFP activity in the mPFC and dCA1 reaching positive peak values significantly above baseline levels for correct but not for incorrect decisions, resulting in a significant decision-outcome related difference in average LFP activity at around 0.4 sec prior to decision completion (Fig. 3C). Photoactivation of LS Purkinje cells eliminated the positive peak in average LFP activity in both mPFC and dCA1 and eliminated the decision-outcome related difference in average LFP activity for correct and incorrect decisions (Fig. 3D).

Decision-outcome related differences in the coherence of mPFC-dCA1 LFP oscillations were observed in four frequency bands in control trials (beta, low gamma, high gamma and high frequency) (Fig. 4A) and these differences were eliminated by LS stimulation (Fig. 4B). Decision-related coherence maxima and minima that occurred in frequency bands between 30 and 200Hz during control trials were also eliminated by LS stimulation (Fig 4). Decision-related coherence in the theta and alpha bands showed no clear differences between control and LS stimulation trials. These findings suggest that Purkinje cells activity in LS plays a crucial role in the modulation of coherence in the beta and gamma bands during SWM decision-making in the mPFC and dCA1.

Theta coherence was seen to increase during decision making, but we did not observe differences in coherence increase between correct and incorrect decisions, which contrasts with two previous studies which reported higher theta coherence values for correct compared to incorrect decisions (Hyman *et al*., 2010; Jones and Wilson, 2005). We suggest that this may be indicative of a specific role of theta oscillations in the decision processes in tasks requiring rule learning compared to decision-making in a spontaneous exploration task. Previous studies used tasks that required some form of rule learning, such as a spatial alternation (Jones and Wilson, 2005), attentional set shift (Benchenane *et al*., 2010) or delayed-non-match-to-sample task (Hyman *et al*., 2010; Sigurdsson *et al*., 2010). These studies not only showed mPFC-hippocampal theta coherence increases during the decision process (4-12Hz), but also an increase in baseline theta coherence correlated with improved task performance (Benchenane *et al*., 2010; Hyman *et al*., 2010; Jones and Wilson, 2005; Sigurdsson *et al*., 2010). A similar increase in theta coherence between the mPFC and hippocampus in correlation with progress in learning was also observed during fear learning (Boyce et al., 2016; Popa et al., 2010). These findings suggest that theta oscillations and theta coherence may be involved in memory consolidation (Boyce *et al*., 2016) which is required for rule learning but not for spontaneous behavior. Our results show an involvement of coherence in multiple higher frequency bands in the decision process during spontaneous exploration. Previous studies have not reported results from higher frequencies which leaves open the question of whether coherence changes in higher frequencies did occur in rule-learning based decision processes. To address this question experiments comparing both types of decision-making in the same animals would be needed.

The effects of optogenetic activation of LS Purkinje cells on SWM behavioral outcomes and the modulation of mPFC-dCA1 coherence during SWM based decision-making suggest a causal involvement of the cerebellar LS in SWM decision making and mPFC-dCA1 beta and gamma coherence modulation. We propose that the cerebellum contributes to the cognitive process of SWM decision-making by optimizing the task-related modulation of mPFC-dCA1 gamma coherence. A proposed role of the cerebellum in optimizing task related coherence between frontal cortical areas and hence optimizing neuronal communication in a task related manner is also supported by experimental evidence in the sensorimotor domain (Lindeman et al., 2021; Popa et al., 2013). Popa et al. showed that inhibiting cerebellar output from the interposed nuclei in awake, head fixed mice reduced gamma coherence between sensory and motor cortical areas (Popa *et al*., 2013). Lindeman and colleagues demonstrated cerebellar involvement in a task related modulation of theta and gamma coherence between sensory and motor areas involved in active whisker movements (Lindeman *et al*., 2021).

Cerebellar involvement in the coordination of cerebral-cortical coherence to optimize task related neuronal communication might thus apply to both cognitive and sensorimotor functions. The coordination of coherence is in essence a temporal coordination problem, as it requires the precise temporal alignment of the phases of oscillations between two structures. The cerebellum has long been recognized as a key structure for the analysis and coordination of precisely timed events, as required, for example, in the control of muscle contractions to optimize motor coordination (Braitenberg et al., 1997; Diener et al., 1992; Ivry and Spencer, 2004; Raghavan et al., 2016). The same principle of precise temporal coordination can here be applied to the phase alignment of neuronal oscillations to optimize coherence and thus cerebral cortical communication in any task dependent manner. Such a role in modulating the communication between cerebral cortical areas would engage the cerebellum in almost any task involving the cerebral cortex and could explain why functional imaging studies reliably detect activation of the cerebellum in a broad range of sensorimotor and cognitive tasks (Zacharia and Eslinger, 2019). Our findings in the cognitive domain and those of others in sensorimotor context strongly suggest the possibility that the cerebellum has a role in coordinating communication through coherence between cerebral cortical areas. Future studies of cerebrocerebellar interactions should take this possibility into account by monitoring changes in functional connectivity between cerebral cortical areas and relate them to changes in activity in the cerebellum.

## MATERIALS AND METHODS

### Animals

Nine adult Tg(Pcp2-COP4*H134R/EYFP) U126Isop/J mice (4 female, 5 male), co-expressing Channelrhodopsin2-EYFP in cerebellar Purkinje cells were used in this study. Mice were housed in a climate-controlled mouse colony at the University of Tennessee Health Science Center animal facilities with 12-hour light/dark cycles in standard cages with free access to food and water. All animal procedures were performed in accordance with the NIH Guide for the Care and Use of Laboratory Animals (2011). Experimental protocols were approved by the Institutional Animal Care and Use Committee at the University of Tennessee Health Science Center.

### Surgery

Mice were surgically prepared for freely moving electrophysiological recordings in the mPFC and dCA1 (Fig. 2). Surgical anesthesia was initiated by exposing mice to 3% isoflurane in oxygen in an induction chamber. Anesthesia was maintained with 1-2% isoflurane in oxygen during surgery using an Ohio isoflurane vaporizer (Highland Medical Equipment, Deerfield, IL, USA). Body temperature was maintained at 37-38°C with a servo-controlled heat blanket (FHC, Bowdoinham, ME, USA) monitored by rectal thermometer. At the beginning of each surgery, after mice were anesthetized but before the first incision, mice received a single subcutaneous injection of the analgesic Meloxicam SR (4 mg/kg, 0.06 ml) to alleviate pain. Two round openings (1.0 mm diameter) were prepared in the skull bone overlying the left mPFC (AP 2.46 mm; ML 0.5 mm) and the left hippocampus (AP −2.3 mm; ML 2.0 mm) (Fig. 2A) using a dental drill (Microtorque II, RAM Products, Inc., USA), leaving the underlying dura intact. A third small opening was prepared overlying the LS and a 5 mm long optical fiber (200 micrometers diameter, Doric Lenses, Canada) coupled to and LED (465 nm, 17 Watts, Doric Lenses, Canada) which was fixed in place with acrylic cement (Co-Oral-Ite Manufacturing, USA) so that the fiber would touch but not penetrate the dura overlying LS. The LED was powered via a thin wire (2 m, 40 AWG solid nickel) connected to two miniature female gold plugs, which were also embedded in acrylic cement.

Two extracellular recording electrodes (glass insulated tungsten/platinum; 80 μm diameter; impedance: 3.5-5.0 MΩ) attached to a custom-made micro-drive were centered over the mPFC and dCA1 skull openings and the micro-drives were fixed to the skull using acrylic cement. The four electrodes, a reference wire and a ground wire were then connected to a 20-pin micro-connector (Omnetics Connector Corp.). Finally, an acrylic head-post was mounted on the skull to provide a handle to manually stabilize the head while connecting and disconnecting the wireless head stage. The micro-drives, head-post and micro connector were embedded in dental cement and anchored to the skull bone using three small skull screws. Of those three screws, one on the right side (AP −1 mm; ML 3 mm) was connected with the reference wire and one on the left side (AP −4 mm; ML 4 mm) was connected to ground. A postsurgical recovery period of 3-4 days was allowed before electrophysiological experiments began.

### Electrophysiology

Electrophysiological recordings were conducted with extracellular recording electrodes (glass insulated tungsten/platinum; 80 μm diameter; impedance: 3.5-5.0 MΩ, Thomas Recording, GmbH, Germany) attached to a custom-made micro-drive. Two electrodes, separated by 0.25 mm, were placed in the mPFC and dCA1. All four electrodes were connected to a 20-pin micro-connector (Omnetics Connector Corporation). During recording sessions, a wireless head stage (W2100-HS16, Multichannel Systems, Germany) was plugged into the micro-connector. To reduce weight the battery was kept off the head stage and power was supplied by connecting battery and head stage via two highly flexible thin wires (2 m, 40 AWG solid nickel). Electrophysiological recordings were performed on five consecutive days. On the first day, electrodes were manually advanced into the left mPFC and dCA1 while the animals were in their home-cage. The occurrence of sharp wave ripples (SWR) were used to determine electrode tip placement in dCA1 (Buzsaki, 2015) (Fig. 2B). On subsequent days, recordings were performed continually while the mice freely explored the plus maze for 12 min. Electrode positions were only altered in the dCA1 if SWR signals were lost. Broad band voltage signals (0.1-8 kHz) were digitized at 20 kHz and saved to hard-disk (W2100-HS16, Multichannel Systems, Germany). LFPs were band-pass filtered off-line at 0.1-200 Hz and down-sampled to 2 kHz using Spike2 software (Cambridge Electronic Design Limited, UK). After completion of the final experiment, recording sites were marked by small electrolytic lesions (10 μA DC; 12 s) and verified anatomically (Fig. 2C).

### Behavioral task

The plus-maze task was used to quantify SWM using counts of spontaneous alternations (Richman *et al*., 1987) (Fig. 1). During spontaneous maze exploration, healthy mice tend to avoid entering recently visited arms, resulting in an above-chance occurrence of arm-entry sequences without repetition. Such repeat-free sequences are called *spontaneous alternations* (Richman *et al*., 1987) (Fig. 1B). The sequence of arm entries is automatically extracted from the video recordings of maze exploration behavior, with arms represented by a letter (A,B,C,D) and analyzed in accordance with established procedures (Richman *et al*., 1987). In creating the sequence of entries, re-entries into the same arm are ignored. Thus, if the mouse does not choose a new arm but returns into the current arm, the letter representing the current arm occurs only once in the sequence. The percentage of spontaneous alternations are calculated by searching the sequence of arm entries for sequences of four entries (i.e. four letters) without a repeat. To this end, a four-letter-wide sliding window is moved along the sequence, advancing the window one letter at a time. Spontaneous alternations can be overlapping, as shown in Figure 1B. The percentage of spontaneous alternations is calculated by dividing the number of no-repeat four-letter-sequences by the total number of four-letter-sequences, multiplied by 100 (Richman *et al*., 1987). Spontaneous alternation percentages are then averaged across mice and compared between control and the LS stimulation trials and also compared to chance level performance. The number of spontaneous alternations expected by chance were evaluated by generating a random sequence for each mouse (n=9) with 50 arm visits in each random sequence.

The random values for the Plus-task performance were generated in Matlab based on the rules for calculating spontaneous alternations. To match the sample size and the average number of entries per task performance, nine randomized sequences of data were created, each contained 50 choices. Two-Way ANOVA were used to compare SWM performance (spontaneous alternation percentage) between control, LS stimulation and random plus maze trials.

These random sequences were then analyzed the same way as observed sequences of arm entries, revealing an estimated chance level performance of 22.2% spontaneous alternations. Healthy mice or rats generate significantly higher numbers of spontaneous alternations and a decrease in spontaneous alternations is interpreted as a deficit in SWM (Lalonde, 2002; Richman *et al*., 1987). Mice explored the plus-maze for 12 min while their movement trajectory was automatically tracked with a video system (30 frames/s; Viewer, Biobserve GmbH). Arm entry sequences and resulting spontaneous alternations were analyzed off-line.

Each mouse performed the plus-maze test five times, once per day, on five successive days. The first session was to allow mice to become familiar with the maze and was not analyzed further. On the four subsequent maze test days, electrophysiological recordings were performed during each session and mouse movements were tracked by video. On two out of four test days mice receive optical stimuli to the cerebellar LS at the time of decision-making. We alternated between days with and without optical stimulation. Which mouse received optical stimulation on days 1 and 3 or days 2 and 4 was pseudo-randomized.

In order to link brain activity to decision-making outcome, we classified individual decisions as correct and incorrect, depending on where the decision occurred relative to a spontaneous alternation (Fig. 1C). All decisions outside spontaneous alternations were classified as incorrect. In classifying decisions falling within spontaneous alternations we chose the following conservative approach: of the four decisions that occur within a single spontaneous alternation, we considered the first three decisions to be part of a sequence of four that includes at least one repetition (Fig. 1B, SA1) and thus classified them as incorrect. Only the fourth decision in the sequence was classified as correct. If multiple spontaneous alternations occur uninterruptedly every decision after the third was classified as correct (Fig. 1B, SA 2-5).

### Optical stimulation of LS PCs

Optical stimulation was performed with a fiber-coupled LED (465 nm; 17 mW, Doric Lenses, Canada) mounted on a 5 mm long optical fiber (200 micrometers diameter, Doric Lenses, Canada), with the optical fiber placed directly on the cortical surface of the LS without penetrating the dura. This method generated no significant electrical artifacts in LFP recordings and allowed the analysis of mPFC-dCA1 coherence during trials with optical stimulation.

LS photo stimulation was applied during the beginning of the decision-making process. Stimulus onset was manually timed to occur at the moment the mouse reached the center area of the plus maze (blue shaded area in Fig. 3A, right panel). Photo stimuli consisted of a 1 sec light stimulus (sinusoidally modulated illumination, 120 Hz). Sinusoidal modulation of light was chosen to avoid the possibility of a depolarization block to occur with sustained DC illumination. The sinewave voltage controlling LED illumination was digitized at 2 kHz and recorded to hard disc together with the electrophysiological data.

### Raw data processing

Raw electrophysiological data were first processed to remove power-line interference (60 Hz and harmonics) using a spectrum interpolation method (Leske and Dalal, 2019; Mewett et al., 2004). Data were then low pass filtered to create LFPs for further analysis (cutoff frequency: 200 Hz). All LFP signals were aligned at the time point that marked the moment the mice left the center area with all four paws, and hence had completed the decision process. This time (time 0 in all decision time-aligned plots) was determined by analyzing the video offline. All decision-aligned data analysis of average LFP activity and coherence was performed on eight-second-long segments of LFP data centered around time 0 for each decision.

### Time-resolved coherence analysis of LFP

Of the two LFP signals available from two recording sites in each the mPFC and in the dCA1, one signal was chosen for further analysis. In the dCA1 we chose the LFP with the highest amplitude SWRs for further analysis, unless there was no difference, in which case one signal was chosen at random. In the mPFC, if LFP signals were equal in quality and one was chosen at random. In both the dCA1 and the mPFC recordings, if one signal showed increased line noise or artifacts, the other was used. All LFP data were first z-scored (Matlab function code: zscore). Time-resolved coherence was calculated in Matlab using custom scripts (Matlab R2019b; function code: *wcoherence*).

### Statistical analyses

Two-Way ANOVA were used to compare SWM behavioral performance (spontaneous alternation percentage) between control, LS stimulation and random plus maze trials.

The peri-decision LFP and coherence time courses were analyzed for each individual decision, separating correct and incorrect decisions (Fig. 1). To allow between-mouse comparison of LFP signals, all signals were z-scored. LFP trajectories for correct and incorrect decisions were compared within 50 ms wide sliding windows (Fig. 3C,D). Coherence trajectories for correct and incorrect decisions were calculated for each frequency range separately. As a first step baseline coherence values were subtracted from each point of the peri-decision coherence function, with baseline coherence defined as the average coherence in the 0.5 s period immediately after the mouse entered the center square of the maze with all for paws. Note the time point of center entry has a variable relationship with the time of completion of decision process (t = 0 in the peri-decision coherence plots in Fig. 4) and typically occurs about 3 s prior to t=0.

Next maxima and minima in the peri-decision coherence time courses were detected. Wilcoxson Signed-Rank test was used to determine whether coherence maxima or minima for correct or incorrect decisions reached values that were significantly higher or lower than the pre-decision baseline coherence values. Open green and red triangles in Fig. 4 mark coherence peaks for correct and incorrect decisions, respectively, that exceed baseline coherence averages (Wilcoxon Signed-Rank test). Similarly, red open circles mark coherence minima falling below baseline coherence values for incorrect decisions (Wilcoxon Signed-Rank test). No significant coherence minima were detected for correct decisions; hence no green open circles were placed. To test for differences between coherence trajectories for correct and in correct decisions, baseline-subtracted coherence values were averaged within sliding windows with window-width equal to one time period of the lowest frequency in the range (Fig. 4). This lowest frequency span-based time resolution preserved the contributions of the entire frequency range in the time-frequency averaged spectral quantities (Mitra and Pesaran, 1999). Decision-outcome dependent differences in LFP amplitude and coherence were evaluated using Two-Sample t-tests. Figures represent results as mean ±standard error.

### Histological evaluation of recording location

At the end of the experiments, animals were deeply anesthetized with intraperitoneal injection of Avertin (Tribromoethanol, 500 mg/kg) and intracardially perfused with 0.9% NaCl and followed by 4% paraformaldehyde solution. Brains were removed from the skull and post-fixed in 4% paraformaldehyde solution for a minimum of 24 hours. Fixed brains were then sectioned at 60 μm, mounted onto glass slides and Nissl stained. Light microscopy was used to localize electrolytic lesions and verify the correct placement of the recording electrode tip in the mPFC and the dCA1 (Fig. 2C).

## ACKNOWLEDGEMENTS

We would like to thank the Neuroscience Institute of the University of Tennessee Health Science Center (UTHSC) for financial support, Micheal Nguyen from the UTHSC Bio-Medical Services and Shuhua Qi for technical support. We would also like to thank Amanda Brown and Trace Stay (Baylor College of Medicine) for support with some of the breeding and also for genotyping.

## AUTHOR CONTRIBUTIONS

Conceptualization: D.H.H., S.S.M, R.V.S., Y.L.; Formal Analysis: Y.L., S.S.M. and M.D.; Funding Acquisition: D.H.H and R.V.S.; Methodology: Y.L.; Supervision: D.H.H.; Writing – original draft: Y.L. and D.H.H.; Writing – review & editing: D.H.H., Y.L., S.S.M, M.E.v.d.H, M.D., R.V.S.

